# Pharmacological prevention of neonatal opioid withdrawal in a pregnant guinea pig model

**DOI:** 10.1101/2020.07.25.221192

**Authors:** Alireza Safa, Allison R. Lau, Sydney Aten, Karl Schilling, Karen L. Bales, Victoria A. Miller, Julie Fitzgerald, Min Chen, Kasey Hill, Kyle Dzwigalski, Karl Obrietan, Mitch A. Phelps, Wolfgang Sadee, John Oberdick

**Affiliations:** Department of Neuroscience, The Ohio State University Wexner Medical Center, Columbus, OH; Department of Psychology, California National Primate Research Center, Animal Behavior Graduate Group, University of California, Davis; Anatomisches Institut, Anatomie und Zellbiologie, Rheinische Friedrich-Wilhelms-Universität, Bonn, Germany; Division of Pharmaceutics and Pharmaceutical Chemistry, College of Pharmacy, The Ohio State University, Columbus, Ohio; Dept of Cancer Biology and Genetics, Ohio State University Wexner Medical Center, Columbus, OH; Aether Therapeutics Inc., 4200 Marathon Blvd. Austin, TX 78756

**Author notes:** To whom correspondence should be addressed (J.O.) and (W.S.). These authors contributed equally to the manuscript (A.S., A.R.L., and S.A.). Email addresses of all authors (A.S.). (A.R.L). (K.S.). (K.L.B.). (S.A.). (V.A.M.). (J.F.). (M.C.). (K.H.). (K.D.). (K.O.). (M.A.P.). (W.S.). (J.O.).

**Keywords:** opioid, neonatal, withdrawal, guinea pig, preventive, therapeutic, HPA axis, cortisol

## Abstract

Newborns exposed to prenatal opioids often experience intense postnatal withdrawal after cessation of the opioid, called neonatal opioid withdrawal syndrome (NOWS), with limited pre- and postnatal therapeutic options available. In a prior study in pregnant mice we demonstrated that the peripherally selective neutral opioid antagonist, 6β-naltrexol (6BN), is a promising drug candidate for preventive prenatal treatment of NOWS. Here, we have developed methadone (MTD) treated pregnant guinea pigs as a physiologically more suitable model, enabling detection of robust spontaneous neonatal withdrawal. Prenatal MTD significantly aggravates two classic maternal separation stress behaviors in newborn guinea pigs: calling (vocalizing) and searching (locomotion) - natural attachment behaviors thought to be controlled by the endogenous opioid system. In addition, prenatal MTD significantly increases the levels of plasma cortisol in newborns, showing that cessation of MTD at birth engages the hypothalamic-pituitary-adrenal (HPA) axis. We find that co-administration of 6BN with MTD prevents these withdrawal symptoms in newborn pups with extreme potency (ID50 ∼0.02 mg/kg), at doses unlikely to induce maternal or fetal withdrawal or to interfere with opioid antinociception based on many prior studies. Furthermore, we demonstrate a similarly high potency of 6BN in preventing opioid withdrawal in adult guinea pigs (ID50 = 0.01 mg/kg). This suggests a novel receptor mechanism to account for the selectively high potency of 6BN to suppress opioid dependence as compared to its low potency as a classical opioid antagonist. In conclusion, 6BN is an attractive compound for development of a preventive therapy for NOWS.

## INTRODUCTION

Extended prenatal exposure to an opioid, with cessation at birth, may result in neonatal opioid withdrawal syndrome (NOWS) *(1)*. NOWS babies are often born premature and underweight, are typically extremely irritable with inconsolable high-pitched crying, with uncontrolled and jittery limb movements, disrupted sleep, and other complications. These problems result in extended ICU stay times, accounting for most of the financial costs of NOWS *(2,3)*. In addition, there are significant later-life effects, including motor and cognitive delay, with untold long-term costs *(4,5)*. Opioids continue to be the best relief for chronic or severe pain, and as many as 28% of pregnant women are reported to have filled an opioid prescription for pain or opioid use disorder management *(1)*. Buprenorphine, a favored agonist for maintenance therapy, results in less severe neonatal withdrawal than methadone (MTD), but both agonists have a similar rate of NOWS after birth (∼50%), and both require significant ICU stay times *(50)*. Here we evaluate a prenatal preventive strategy that, in principle, should reduce long ICU stay-times, the need for postnatal morphine treatment of NOWS babies, and the negative developmental consequences of prenatal opioids. Currently only palliative therapies are available to those babies that develop NOWS, and none would prevent developmental delay.

In a previous study we demonstrated the preferential delivery of a neutral mu-opioid receptor antagonist, 6β-naltrexol (6BN), to the fetal brain in pregnant mice *(6)*. While partially excluded from the maternal brain, 6BN rapidly transits the placenta and enters the fetal brain reaching levels 6-fold higher than in maternal brain. The relative exclusion of 6BN from maternal brain is thought to be due to efflux transporters at the mature blood brain barrier (BBB), such as P-gp *(7)*. The developmentally regulated expression of such transporters at the BBB may account for the observation that 6BN readily entered the fetal brain and brains of pre-weaning juveniles until at least postnatal day (PD) 15 *(6)*. Inducing morphine dependence in juvenile mice, we demonstrated that co-administration of 6BN at extremely low doses prevents subsequent withdrawal behavior with a 50% inhibitory dose (ID50) of 20-40 ug/kg – 500-fold lower than the dose of morphine used to induce dependence in the study, and 50-100-fold lower than the published ID50 of 6BN to block opiate antinociception or to induce withdrawal in opioid dependent adult animals, including in mice and monkeys *(8-13)*. Here, we test the hypothesis that 6BN can prevent fetal dependence and subsequent neonatal withdrawal when co-administered at extremely low doses with an opioid agonist – at 6BN doses too low to interfere with maternal pain or addiction management by an opioid.

Facile 6BN access to the immature mouse brain *(6)* should account for no more than 5-10-fold higher potency in blocking opioid antinociception compared to adults with an intact BBB. Therefore, it is likely that factors other than the BBB are at play, and alternative mechanisms at the opioid receptor likely play a role in 6BN’s extremely high potency to prevent dependence in mouse juveniles as compared to its low potency in blocking other agonist actions in adults. We address this question here by also testing the ability of 6BN to prevent naloxone induced withdrawal when co-delivered with MTD to induce dependence in adult guinea pigs.

There are some limitations of rodents as a model for NOWS. Mice show no reported behavioral effects of prenatal MTD at birth, but rat pups display spontaneous and naloxone-induced increased movements *(14-16)*. The latter behaviors are strictly count-based binary measures (yes-no) of limb, body and head moves. The subtlety of these behaviors is presumably due to the relative underdevelopment of the brain of these species at birth, which are the neurodevelopmental equivalent of mid-second trimester human fetuses *(17)*. However, more robust withdrawal behaviors in the locomotor and ultrasonic vocalization domains can be observed by PD7-10 in rat pups using postnatal exposures to an opioid *(6,18)*. Mice and rats at this age are the neurodevelopmental equivalent of human newborns and may be useful for modeling some aspects of NOWS. But they lack the temporal continuity of opioid cessation at birth, as occurs in NOWS.

In order to develop a more human-relevant model we have been studying guinea pigs, a precocial species developmentally more akin at birth to a human infant *(17)*. These studies suggest that cessation of an exogenous opioid at birth after extended prenatal exposure causes an increased “craving” for the opioidergic effects provided by infant-parent contact, resulting in NOWS, and likely engaging stress mechanisms mediated by the hypothalamic-pituitary-adrenal (HPA) axis.

Finally, we show for the first time that 6BN can prevent spontaneous withdrawal behaviors after birth when co-administered with a prenatal opioid, and it can do so with high potency. Moreover, the high potency does not appear to require preferential delivery of 6BN to the fetal CNS.

## MATERIALS AND METHODS

### Animals

Hartley guinea pigs (Cavea porcellus) were purchased from Charles River. All adult non-pregnant animals were 2-4 mo old at the time of testing. Pregnant sows for PK studies were purchased in 3 cohorts of 7 animals each. Pregnant sows for neonatal behavior studies were purchased in 6 cohorts of 8 – 10 animals per cohort. All pregnant animals were shipped at early-gestation (∼GD30) to reduce pregnancy-related toxemia, which is a significant concern for guinea pigs. The standard range of pup birthweights for Hartley guinea pigs is 50 - 100 g *(51)*. However, for in-house reared guinea pigs, birthweights below 85 g are sometimes considered low birthweight, and “normal” birthweight is considered those on the high end of the normal range, >90 g *(52)*. Thus, the relatively lower mean birthweight of 75.5±3.1 g for saline control animals in the current study (Tables 5 & 7) may reflect vendor-related substrain differences, but we cannot rule out an impact of shipping-related stress.

Pregnant sows were typically second or third pregnancies, and 6 to 8 month old. These were “multi untimed pregnant animals” with a 3 day window of variation for day of conception (gestation time of guinea pigs is ∼65 days). Animals were pair-housed whenever possible in solid bottom open caging (Allentown, Allentown, NJ, cage pans measured 29 × 21 x 10 inches) with Sani Chip bedding (Teklad 7090, Envigo). Animals were fed a chow diet ad libitum (Teklad Guinea Pig Diet 2040, Envigo) that was supplemented with hay and a rotation of fresh produce provided daily. Reverse osmosis purified water was provided via an automated rack watering system, and a 12/12 hour light/dark cycle was maintained for the duration of the experiment.

All procedures were approved by The Ohio State University Institutional Animal Care and Use Committee and are in compliance with guidelines established by the National Institutes of Health published in Guide for the Care and Use of Laboratory Animals (http://oacu.od.nih.gov/regs/guide/guide.pdf).

### Drugs

6β-naltrexol (6BN) (https://pubchem.ncbi.nlm.nih.gov/compound/5486554), naltrexone (https://pubchem.ncbi.nlm.nih.gov/compound/5360515) and d,l-methadone (MTD) (https://pubchem.ncbi.nlm.nih.gov/compound/14184) were provided by the Drug Supply Program of the National Institute for Drug Addiction (NIDA) as previously reported *(6)*. Drugs were dissolved in saline at concentrations between 10 and 20 mg/ml, and all dilutions were made in saline.

### Pharmacokinetic (PK) analysis

#### 6BN and naltrexone dosing and tissue collection

For the bioavailability study young adult (2 mo old) animals with jugular vein catheters were purchased from Charles River for plasma multi-sampling and IV dosing. Half of the animals were used for IV, half for oral delivery. For oral delivery animals were briefly anesthetized with isoflurane, held vertically, and drug (in saline + 1% sucrose) was delivered with a feeding needle, typically in 1.5 - 3 ml. This procedure induced significant salivation and/or regurgitation, which contributed to dosing variability. For the PK studies in brain and plasma most of the reported data used oral delivery.

However, as indicated in the Results other animals were injected subcutaneously in the dorsolateral region around the shoulders of the forelimbs. Injection volume typically did not exceed 750 ul in adults. Embryos and maternal tissues were collected for the plasma and brain PK study. After drug injection and variable survival times pregnant animals were euthanized by CO2 followed by heart puncture (for maternal blood collection, ∼0.5 – 1 ml) and decapitation. Fetuses were collected onto large petri dish lids kept on wet ice, and blood was collected in a 1 ml syringe (no needle) from pooling of blood around the neck region after severing of both jugular veins (with animal on its back). Then brain and liver were collected and frozen on dry-ice while plasma was prepared in a microfuge at room temp (typically from 200-400 ul of fetal blood). All samples were stored at -70oC until processing. In the mass spectrometry lab tissues were thawed, resected, weighed, and quickly frozen again on dry ice in individual microcentrifuge tubes. Samples were later thawed, processed and analyzed via liquid chromatography-tandem mass spectrometry (LC-MS/MS) to quantify levels of drug.

#### Bioanalytical Assay for 6β-naltrexol and naltrexone

Guinea pig plasma and brain tissue were assayed according to Oberdick, et al (PMID: 27189967) (6) with the following changes going from mouse to guinea pig plasma and tissues. Guinea pig brain and liver tissues were homogenized in 1X PBS pH 7.4 at concentrations of 66.7 mg/mL and 33.3 mg/mL, respectively. Forty-five microliters of plasma or tissue homogenate were spiked with 5 µL of 3000 ng/mL [^2^H-7]-naltrexone or a combination of [^2^H-7]-naltrexone and 6β-naltrexol-d3. Samples were then extracted with acetonitrile:methanol (3:1, v/v). The post centrifugation supernatant was transferred to a new 2 mL 96-well plate then dried with a stream of nitrogen before reconstituting with 120 µL of 0.1% formic acid. The reconstituted samples were transferred to a 96-well autosampler plate for a 5 µL injection onto a Thermo Accucore Vanquish C18+ column (1.5 µm, 100 × 2.1 mm). The chromatographic system was a Thermo Scientific Vanquish Horizon UHPLC. The analytes were separated using a 3.5 minute gradient elution program with 0.1% formic acid (MPA) and acetonitrile:methanol (3:1, v/v) with 0.1% formic acid (MPB) as the mobile phases. The gradient began with 10% MPB for 0.4 minutes then increased to 90% MPB over 1 minute. Then 90% MPB was held for 0.6 minutes before decreasing back to 10% MPB to equilibrate the column. The flow rate for the gradient was 0.4 mL/min and the column was maintained at 40 °C. The samples were analyzed on either a Thermo Scientific TSQ Quantiva or a TSQ Altis equipped with an electrospray ionization source in positive polarity. The mass transitions (m/z) monitored were 342.18 > 324.11 for naltrexone, 344.21 > 326.11 for 6β-naltrexol, 349.24 > 331.11 for [^2^H-7]-naltrexone, and 347.36 > 329.25 for 6β-naltrexol-d3. Cross-validation included linearity, within-day, and between-day accuracy and precision. Selectivity was assessed by analyzing six lots of guinea pig plasma without and with spiking at the lower limit of quantitation (LLOQ) levels. Recovery, matrix effects and stability were also assessed in matrix and solvent for both analytes at low and high QC concentrations. Stability was assessed on benchtop (22 °C), three freeze thaw cycles from -80 °C to 22 °C, autosampler stability for 24 hours at 4 °C, and long-term stability at -80 °C for 30 days. The linear range, including LLOQ for 6BN and naltrexone were as follows: Plasma, x-y ng/mL 6BN, x-y ng/mL naltrexone; Liver, x-y ng/mL 6BN, x-y ng/mL naltrexone; Brain, x-y ng/mL 6BN, x-y ng/mL naltrexone.

### Pharmacokinetic Data Analysis

For the bioavailability study, non-pregnant adult guinea pigs were dosed with 6BN via intravenous bolus route at 10 mg/kg and orally at 40 mg/kg. Plasma samples were collected at 0, 15, 30, 60, 120, 240 and 480 min post dose. Individual 6BN pharmacokinetic parameters were estimated from individual plasma concentration-time profiles using non-compartmental analysis (NCA) with Phoenix WinNonLin (version 8.2.0, Certara, Princeton, NJ). The terminal linear phase was identified automatically in WinNonlin using linear least squares regression to estimate the terminal elimination rate constant (λz). Area under curve (AUC) was determined using linear trapezoidal linear interpolation method. Bioavailability was calculated as the dose-normalized ratio of AUCs determined from oral and intravenous routes of administration. For the brain distribution analysis, the concentration-time profile for both brain and plasma were anlyzed with NCA methods using Phoenix WinNonLin and PK parameters were calculated in a similar fashion. Pregnant guinea pigs were dosed with 6BN orally at 40 mg/kg, then both fetal and maternal brain and plasma samples were collected at 15, 30, 60, 120, 240 and 480 min post dose (one time point per animal).

### Protein binding by equilibrium dialysis

Data were collected and analyzed by Covance.

#### Instrument

The high throughput dialysis apparatus, model HTD96b, was used (HTDialysis LLC, Gales Ferry, Connecticut). Prior to assembly, dialysis membrane strips (molecular weight cutoff of 6 to 8 kDa) were hydrated according to manufacturer’s recommendations. The Teflon bars were assembled according to manufacturer’s instructions, with dialysis membrane strips laid between bars creating two compartments per well. The assembled unit was locked in place in the steel base plate. Samples were immediately added to each compartment to prevent dehydration of the membranes.

#### Equilibrium Dialysis Procedure

Fortified matrix (plasma) was added to the donor side, DPBS was added to the receiver side of the HTD wells, and the plate was sealed. Samples were incubated at 37°C and rotated at 300 rpm for the designated time. After incubation, the seal was removed and a sample from each plasma and dialysate chamber was analyzed by LC-MS/MS. All protein binding determinations were performed in quadruplicate. *Time to Equilibrium Determination*. Dialysis was performed according to the equilibrium dialysis procedure for 3, 5, 6, 7, and 8 hours to determine the minimum time to achieve equilibrium. This experiment was conducted at 50 ng/mL of 6-beta naltrexol in human plasma. Equilibrium was obtained when the percentage of 6-beta naltrexol bound to the proteins in the plasma remained constant over time.

#### Concentration Dependence

The protein binding in mouse, rat, guinea pig, dog, and human plasma was determined at concentrations of 0.5, 1.5, 5, 15, and 50 ng/mL of 6-beta naltrexol. The dialysis time for the test article was 8 hours as determined in the time to equilibrium experiment. *Sample Analysis*. Samples were processed prior to LC-MS/MS analysis as follows. The donor side samples (plasma) were diluted with DPBS and the receiver side samples (DPBS) were diluted with blank control plasma at the appropriate volumes to provide a common analytical mixed matrix of 90% DPBS and 10% plasma (90:10 DPBS:plasma, v:v) in 100 µL total volume. Samples were prepared for analysis and analyzed for 6-beta naltrexol using a quantitative LC MS/MS method developed at Covance and modified as appropriate for study optimization.

### MTD and 6BN dose-response studies in newborn and adult guinea pigs Experimental design for newborn studies

#### Overview

In this study 10 mg/kg was identified as the maximum tolerable dose of MTD for both sow and fetus survival. Thus, we determined the prenatal MTD dose response for neonatal withdrawal in the range from 0 – 10 mg/kg. Then we selected two MTD doses near the mid-point of the dose-response curve for testing whether 6BN could prevent the neonatal effects of MTD.

#### Dosing groups and animal numbers

All behavioral studies were performed between 12 – 4 PM to ensure similar circadian schedules. Generally, tests were performed in cohorts of 20-30 pups from 8-10 sows at the same gestational stage (with a 3 day window of variation; see Animals section). With sow numbers thus restricted pups were born over the span of 1 week, which allowed us to maintain the 4 hr testing window each day. Each cohort was designed with multiple dosing groups, always including either a saline control group and/or a MTD only (no 6BN) control group. Pregnant sows were pair-housed, and both animals in a pair received the same treatment.

23 pregnant animals were used for the MTD dose response study, and 37 pregnant animals were used for the 6BN dose response study. No more than 3 and an average of 2 pups per litter were used for all withdrawal analysis to minimize litter effects (for all pregnant animals with all treatments the average litter size was 4.7±1.7(±SD)). Pups for the MTD dose study came from at least two litters per dose group with an average of 4 litters per group. Groups for the MTD study were saline controls (“0” MTD), and 2, 5, 7, and 10 mg/kg MTD. For the analysis of the effects of 6BN, data from animals treated with 5 mg/kg MTD (123 pups; 5-7 litters per 6BN dose group) and 7 mg/kg MTD (30 pups, 2 litters per dose of 6BN) were pooled. This approach seems justified since despite the general MTD-dose dependence of behavior that may be discerned if the whole dose range of 0 - 10 mg/kg is considered (Fig. 3), the behavior of animals treated with 5 or 7 mg/kg of MTD was not detectably different, and the same is true of animals treated with 0 or 2 mg/kg MTD (see Figures 2 & 3). We also note that no statistical interactions between MTD and 6BN could be detected in this combined group. Finally, we note that analysis of the effects of 6BN in the animals treated with 5 mg/kg prenatal MTD yielded very similar results to what was observed for the combined 5 and 7 mg/kg MTD animals, with the obvious differences as one may expect due to a smaller group size.

#### Injection schedule

Pregnant animals were received in the vivarium on ∼GD33, acclimated for 17 d, and daily single injections of saline, MTD, or MTD with 6BN at variable doses were initiated at GD50 for an average of 15±3(±SD) injections before birth at ∼GD65. Pups were marked with an indelible marker on the inside of the ear 24 hr after birth. Pups were behaviorally tested (locomotion and vocalization) at 48 hr after birth (see below).

### Adult withdrawal testing

15 females (non-pregnant) and 11 males were used for this study, 2 – 4 months of age. As for the newborns they were tested in the open-field (see below). Vocalization testing was not performed.

### Behavioral tests

#### Open-field locomotor testing

Open field tests were performed on pups 48±12 hr after birth in order to examine their locomotor behavior before and after naloxone administration. One hour before testing the entire home-cage was transported from the housing room into an enclosed holding vestibule with a door to a separate behavior room. Two cameras (Panasonic model SDR-H80 and Sony Handycam model DCR-SR45) were used to record the animals. One camera (Panasonic) was mounted directly over the open field arena on a stand at a fixed height (104 cm) for all studies. The zoom on the camera was adjusted so that the top and bottom borders of the square arena filled the entire height dimension of the camera’s rectangular field of view. This was the main camera used for video and acoustic analysis. The other camera was placed on the ground adjacent to the arena in order to examine detailed facial expressions and behaviors. The arena was 40 by 40 cm in size, and clean wood chip bedding was placed on the floor of the arena (same type as for housing). Animals were removed from the home-cage in the vestibule, brought into the testing room, and weighed immediately before being placed in the open field arena. Once in the arena they were video-recorded for 10 min. Then they were removed from the arena, injected with naloxone (s.c.) at a dose of 20 mg/kg, and immediately placed back in the arena and video-recorded for 30 min. Note that no more than 3 pups per litter were run in the open field test, with an average of 2 pups per litter. Also, at the start of each video an index card with the date and animal ID was placed briefly in the field of view of the camera and recorded. All videos are archived on external hard-drives and are available to collaborators on an online file sharing service.

Locomotor testing of adults was performed in precisely the same manner using a 10 min open-field test prior to naloxone, followed by injection with naloxone, and a 30 min test in the same arena. <u>Video analysis</u>: Each video was imported into the ANY-maze software (version 6.06), labeled by date and animal ID, and data extracted by an observer blinded to animal treatment. In each video the area of the open field chamber was manually defined using a drawing tool, and a distance of known length within this area was used for standardization. The distance travelled by each animal was calculated based on the displacement of the center point of their bodies (the white coat color of animals was easily detected against the darker wood shavings on the floor of the apparatus). Freezing detection was set up using the default values in the software: a minimum freeze duration of 1000 milliseconds, a “freezing on” threshold of 30, and a “freezing off” threshold of 40. Once all the parameters were entered in the software, the videos were run for analysis. Data from each animal was then assigned to its proper dosage group by another researcher for statistical analysis.

#### Vocalization analysis

WAV audio files were extracted from the same videos that were used for automated video analysis, then analyzed using Raven Pro 1.6 Sound Analysis Software (Center for Conservation Bioacoustics, Cornell Lab of Ornithology, Ithaca, NY) by one researcher who was blinded to the dosage groups. Spectrograms with a 512-point (11.6 ms) Hann window (3 dB bandwidth = 124 Hz), with 75% overlap, and a 1,024-point discrete Fourier transform, yielding time and frequency measurement precision of 2.9 ms and 43.1 Hz were generated. Sounds files were not down sampled.

The features estimated were vocalization count and signal-to-noise ratio (SNR). SNR is the amplitude of the vocalization signal above the background noise. In other words, SNR measured how loud the guinea pigs were, controlling for background noise. In order to count the number of vocalizations emitted by each guinea pig pup in the ten-minute observation in an objective manner, we used the Band Limited Energy Detector to select each vocalization. The Band Limited Energy Detector was tailored to detect the vocalizations of infant guinea pigs. The final detector was validated visually and audibly by one observer (ARL). The target signal parameters for our detector were minimum frequency: 390 Hz, maximum frequency: 920 Hz, minimum duration: 0.032 sec, maximum duration: 0.16 sec, and minimum separation: 0.032 sec. We set our SNR ratio parameters to 45% minimum occupancy with an SNR threshold of 6.0 dB above. In order to estimate SNR of the sound file, one ten-minute selection was generated using the Raven Pro selection tables. SNR was automatically estimated by Raven Pro 1.6.

### Statistical procedures for dose-response studies

Summary data are presented as the means +/-S.E.M. unless otherwise indicated. Two-sided t-tests allowing for variance heterogeneity were used to contrast samples if only two groups were to be compared. To compare multiple groups, and to screen for possible dose-response effects, we used the step-down Tukey trend test adjusted for multiplicity (*(54)*; referred to as Tukey trend test; also known as the Tukey-Ciminera-Heyse trend test). Essentially, this comprises a one-way ANOVA with post-hoc Dunnett tests and the simultaneous fitting of linear, logarithmic and ordinal regressions to the data, while controlling for the multiple testing involved. Again, testing was done such as to account for variance heterogeneity between groups. In the newborn withdrawal studies sex was not considered as a factor due to the difficulty of *accurately* evaluating sex. In the adult withdrawal study sex *was* considered as a factor and showed no significant effect or interactions. Dose-response curves (Figures 7 & 8) were fitted using an emax model *(63)*. These procedures were implemented in R (R Core Team, 2020) using the packages car *(55)*, multcomp *(56)*, tukeytrend *(57)*, sandwich *(58)*, and DoseFinding *(59)*.

The different behaviors tested capture distinct, but correlated, measures of a multifaceted response. To combine them into one common measure, we z-transformed (mean = 0; 1SD = 1) the values of each of the four behavioral measures obtained for all animals (irrespective of their treatment). This puts the values of all behavioral measures on the same scale without distorting their distribution. A composite score (“normalized behavior score”) was then calculated for each animal by adding up the transformed values of the distinct behavioral measures. As all measures except freezing time decreased with increasing doses of 6BN, ranks for freezing time were multiplied by -1 (i.e., inverted) before transformation. This approach was motivated by the goal to obtain a more comprehensive and hopefully more robust characterization of behavior than might be achieved by a single endpoint. It was also motivated by the clinical scores constructed from multiple measurements, and the success of multiple-endpoint analyses in clinical studies (e.g., *(60-62)*, to cite but a few).

We add that if data were put on comparable scales by ranking instead of z-transformation, results fully consistent with those reported above were obtained.

### Analysis of birthweight and maternal weight gain

All data for this analysis were extracted from all pups from all litters described above in the MTD and 6BN dose-response studies. Pregnant sows were weighed at the start of dosing, then every 3-4 d, and then every day as they neared term. Pups were weighed at 24 hr and 48 hr after birth. The birthweight data reported here are at 24 hr.

### Blood collection and cortisol measurement

All samples were collected 48±12 hr after birth. Non-survival trunk blood was collected from pups after 1 hr of maternal separation stress. For the 1 hr separation we removed pups from the home-cage and placed them in a holding cage (plastic mouse cage with lid and air filter) in a different room. Pups were then sacrificed by CO2 asphyxiation followed by rapid decapitation in the vivarium necropsy room. Trunk blood was collected (from the site of decapitation) and put into 1.5 mL Eppendorf tubes coated with EDTA (pH 8.0; Invitrogen Corporation Cat No. 15575-038) to prevent coagulation. Tubes were then centrifuged for 10 minutes at 2500 rpm. Plasma was placed into a separate tube and stored at -80.

Plasma cortisol was assayed by enzyme immunoassay (antibody R4866, produced by UC-Davis Endocrinology Laboratory *(see refs 48,49)* at a 1:1000 dilution. Samples were assayed in duplicate and high and low pools were run with each assay. As all samples were run in one assay, there are no inter-assay cv’s to report. The low controls had an intra-assay cv of 3.61%, while the high controls had an intra-assay cv of 3.58%. The assay was chemically validated for guinea pigs by assessing parallelism and quantitative recovery.

## RESULTS

### Pharmacokinetics of 6β-naltrexol in fetal and maternal guinea pig

We first determined the oral bioavailability of 6BN and other PK parameters in non-pregnant adult guinea pigs (Table 1; Figure 1A). 6BN is orally bioavailable with F = 29%, and t1/2 is 1.1 hr for IV delivery, 2.1 hr by oral delivery. For comparison methadone has a bioavailability range of 36-100% in humans with great interindividual variation, and t1/2 of 12 hr in guinea pigs and 15 - 207 hr in humans *(19-21; see also 22)* – the t1/2 of 6BN is 12h in humans *(66)*. In addition, plasma protein binding of 6BN in all species tested is low over a 6BN concentration range from 0.5 – 50 nM (see Table 2; mean unbound fraction±SD = 91±6% in guinea pig, 95±5% in mouse, 92±11 % in dog, and 90±5% in human). For comparison, the published plasma protein binding of methadone is much higher (mean unbound fraction 11-14%; *(21)*).

**Table 1.**
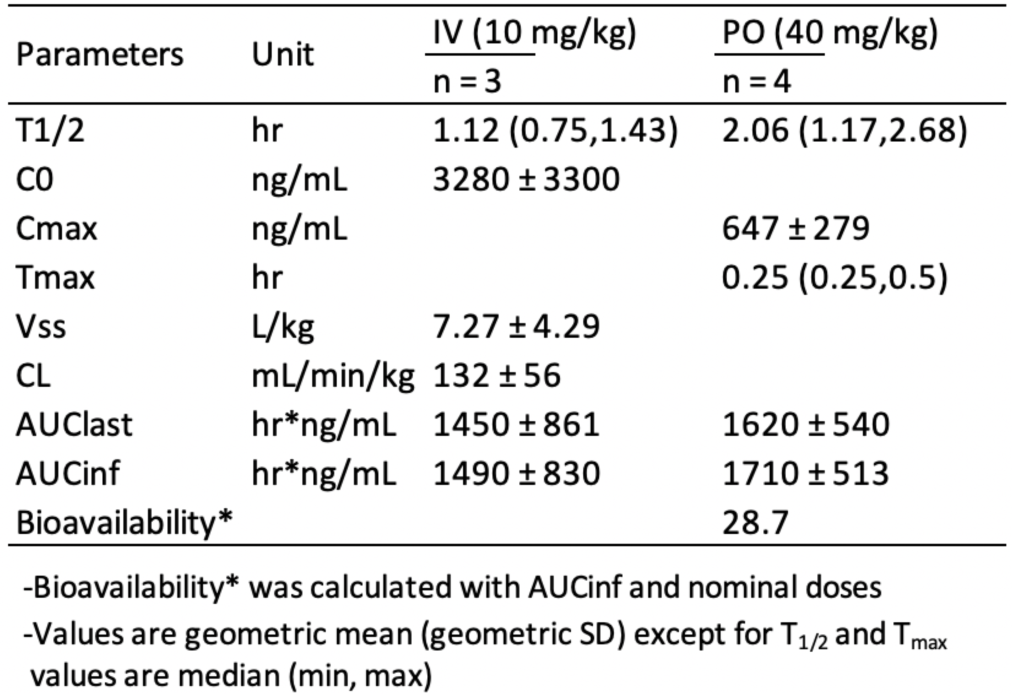
Plasma PK analysis and bioavailability of 6BN

**Table 2.** 6BN mean plasma unbound fraction

| [6BN], nM | Mouse | SD | Rat | SD | Guinea pig | SD | Dog | SD | Human | SD |
| --- | --- | --- | --- | --- | --- | --- | --- | --- | --- | --- |
| 0.5 | 96.5 | 3.36 | 83.5 | 13.10 | 93.7 | 5.53 | 93.2 | 9.40 | 94.3 | 5.22 |
| 1.5 | 96.4 | 3.08 | 90.7 | 7.60 | 82.0 | 3.17 | 90.3 | 11.30 | 90.5 | 6.93 |
| 5 | 90.5 | 6.52 | 91.3 | 4.83 | 94.5 | 8.54 | 94.7 | 6.36 | 95.7 | 5.13 |
| 15 | 97.2 | 4.11 | 87.4 | 10.00 | 89.4 | 8.37 | 90.6 | 13.60 | 77.2 | 0.53 |
| 50 | 92.7 | 5.22 | 92.4 | 9.27 | 95.3 | 5.85 | 90.1 | 12.40 | 93.6 | 7.56 |
| avg | 94.7 |  | 89.1 |  | 91.0 |  | 91.8 |  | 90.3 |  |

data provided by Covance

**Figure 1.**
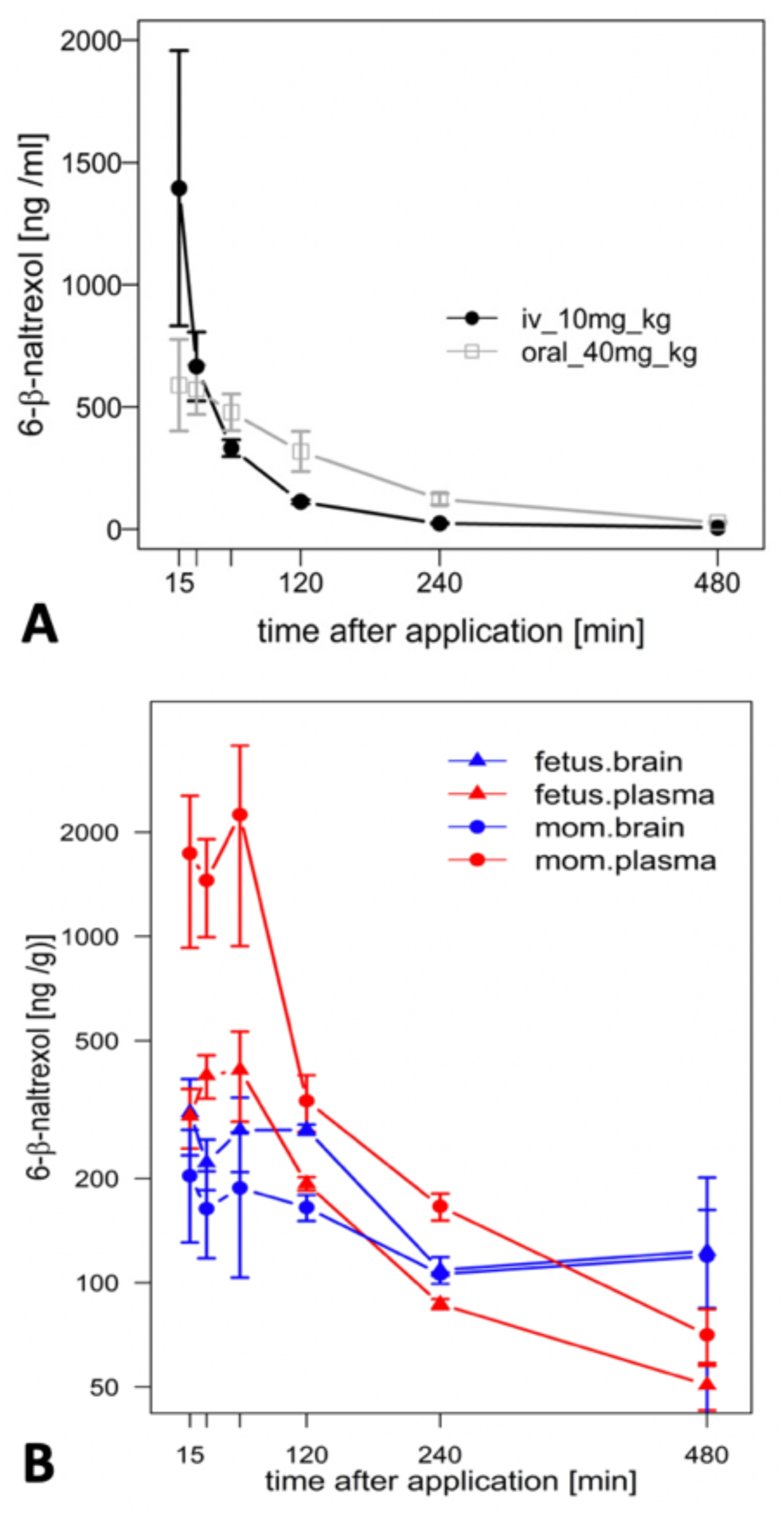
Pharmacokinetic (PK) analysis of 6BN in guinea pigs. A) Bioavailability curves for 6BN in non-pregnant female adult guinea pigs using plasma multi-sampling. The PK time course is shown for oral delivery compared to IV. B) PK time course of 6BN in plasma and brain (dose 40 mg/kg, p.o.). Note y-axis is log scale, in B only. Error bars ±SEM.

We next examined the time course of tissue distribution of 6BN using oral delivery (40 mg/kg; Figure 1B and Table 3). By 1 hr. after dosing 6BN reaches a peak in maternal plasma that is ∼4-fold higher than in fetal plasma, and fetal brain levels are ∼1.5-fold higher than maternal brain levels at 1 – 2 hr. (see AUC_0-4_ in Table 3). In addition, fetal plasma and brain levels are near equivalence at peak in the first 120 min; this result was replicated at a 10-fold lower oral dose of 4 mg/kg 6BN. Using combined data at the two 6BN doses the fetal brain/plasma ratio is 1.04, while the maternal brain/plasma ratio is 0.220 (p < 0.001 for comparison of fetal vs maternal brain/plasma ratio, by t-test). In addition, the absolute level of 6BN in fetal brain in guinea pigs is ∼6-fold lower than in mouse fetal brain under conditions with roughly equal maternal plasma 6BN levels; maternal brain levels are roughly the same across the two species (compare data in Figure 1B to that in *(6)*). These results demonstrate slow placental transfer of 6BN in guinea pigs, but with rapid fetal brain entry, and relative exclusion from maternal brain. Thus, compared to mice, preferential delivery of 6BN to fetal brain is lower in guinea pigs than in mice due to slow placental transfer (i.e., fetal/maternal AUC ratio of brain levels is 1.5 in guinea pig vs 6 in mouse; mouse data reported previously in *(6)*). Due to high variability with oral dosing in pregnant guinea pigs (see Methods) we repeated some of this analysis using subcutaneous delivery with both single injection and multi-injection paradigms. Over a dose range from 0.5 – 10 mg/kg, regardless of the dosing schedule and route, we found slow placental transfer of 6BN in pregnant guinea pigs, similar to that in Fig 1B (data not shown). In addition, while naltrexone showed similar maternal plasma levels as 6BN at the same dose and time after administration, we observed ∼10-fold higher levels of naltrexone compared to 6BN in both fetal and maternal brain (Table S1), indicating more rapid placental transfer and higher maternal brain levels of naltrexone compared to 6BN. These results are consistent with reported relative exclusion of 6BN from adult brain as compared to naltrexone based on pharmacokinetic studies *(6,9)*, and seem to correlate qualitatively, albeit not quantitatively (as discussed in the Introduction), with the greater potency of naltrexone to block opioid analgesia and/or to induce withdrawal in opioid dependent adult animals *(8-13)*.

**Table 3.** PK analysis of 6BN in fetal and maternal plasma and brain

| Parameters | Unit | Mom |  | Fetus |  |
| --- | --- | --- | --- | --- | --- |
|  |  | Plasma | Brain | Plasma | Brain |
| T<sub>1/2</sub> | hr | 2.74 | 11.5 | 3.29 | 6.11 |
| C<sub>max</sub> | ng/mL | 2250 (±1310) | 204 (±73) | 412 (±120) | 310 (±77) |
| T<sub>max</sub> | hr | 1 | 0.25 | 1 | 0.25 |
| AUC<sub>last</sub> | hr*ng/mL | 3810 (±1022) | 1060 (±179) | 1190 (±96.1) | 1360 (±101) |
| AUC<sub>inf</sub> | hr*ng/mL | 4090 | 3040 | 1430 | 2440 |
| AUC<sub>0-4</sub> | h*ng/mL | 3330 | 606 | 910 | 891 |

Values are from same data shown in Figure 1B

### Prenatal methadone aggravates maternal separation stress behaviors in newborn guinea pigs

#### Methadone effect on maternal weight gain and pup birthweight

The standard analgesic dose of MTD for guinea pigs is 3-6 mg/kg *(23)*. However, in a previous study on respiratory effects of prenatal MTD in newborn guinea pigs, 12 mg/kg was found to be the highest dose that did not result in lethality of the pups *(24)*. Therefore, as part of a similar effort to establish the maximum dose of MTD we performed a pilot study on two pregnant animals at this dose. One lost ∼13% of her body weight and was lethargic and unresponsive after 6 days of dosing, and had to be euthanized. The other also lost ∼13% body weight, but over a longer time, gave birth prematurely, and all pups died. We then focused on the prenatal MTD range from 2 mg/kg to 10 mg/kg, first examining maternal weight gain during pregnancy and pup birthweight. As shown in Table 4 control sows gained an average of 206±34 g from the first saline injection to the last injection before birth (average of 15 injections). MTD resulted in a significant dose-dependent decrease in maternal weight gain relative to saline controls with a trend towards reduced weight gain at 2, 5, and 7 mg/kg, no weight gain at 10 mg/kg, and average weight loss of - 175±6 g at 12 mg/kg (Table 4; F(5,18) = 55.7, p < 0.0001, for overall effect of MTD on maternal weight gain, by ANOVA). Also shown in Table 4, pup mortality increased sharply at the 10 and 12 mg/kg MTD dose, but too few animals were analyzed for meaningful statistical analysis. For pup birthweight, those exposed to 5 – 10 mg/kg methadone showed no difference from saline controls, but those at 2 mg/kg had a significantly *higher* birthweight than all others (n = 6 pups from 2 litters; Table 5). Thus, under the conditions of this study MTD had a robust effect on maternal weight gain with little effect on pup birthweight. However, low-dose MTD may lead to increased birthweight, an observation requiring further study since, while significant, there were only 2 litters.

**Table 4.** Effect of methadone on maternal weight gain and litter size

| MTD Dose (mg/kg) | n= | Start weight (g) | End weight (g) | Weight gain (g) | % weight gain | AVG litter size | total pup # | total pup deaths |
| --- | --- | --- | --- | --- | --- | --- | --- | --- |
| 2 | 2 | 1084±40 | 1221±61 | 136±21λ | 12.5±1.5 | 3.0±1.0 | 6 | 0 |
| 5 | 8 | 1195±45 | 1312±64 | 118±27λ | 9.7±2.0 | 5.0±0.4 | 42 | 1 |
| 7 | 2 | 1174±50 | 1295±115 | 121±65λ | 10.1±5.1 | 4.0±1.0 | 8 | 0 |
| 10 | 5 | 1214±28 | 1205±11.9 | -9.2±24.4*** | -0.6±2.0 | 3.8±0.7 | 19 | 7 |
| 12 | 2 | 1335±11 | 1160±5 | -175±6*** | -13.1±0.3 | ND | 5 | 5 |
| 0 (saline ctl) | 5 | 1188±36 | 1395±31 | 206±34 | 17.7±3.4 | 5.8±0.6 | 29 | 3 |

all values ±SEM
F(5,18) = 55.66, p < 0.0001, for overall effect of MTD on maternal weight gain
λF(3,20) = 89.66, p < 0.0001, for overall effect of MTD if data at 2, 5, and 7 mg/kg are pooled as a single group, by ANOVA
\*\*\*p < 0.001 compared to saline controls, by Dunnetts posthoc test
λp = 0.056 compared to saline controls, by Dunnetts posthoc test, if data at 2, 5, and 7 mg/kg MTD are pooled as a single group for ANOVA

#### MTD effect on neonatal withdrawal-related behaviors

To test the effect of prenatal MTD on newborn guinea pig behavior we performed dose-response analysis, testing locomotor and vocalization behavior in an open-field arena (see Methods). The most robust effect of MTD was observed in the 10 min test (before naloxone administration). All newborns, including saline controls, displayed intense spontaneous locomotion immediately upon being placed in the arena. In addition, the pups routinely produced an audible high-pitched call (see Supplemental Movie S1). These are maternal separation behaviors that have been previously described, consisting of an initial active seeking and calling phase, followed by a more protracted “despair” phase *(25-27)*. The effect of MTD was examined at four doses: 2, 5, 7, and 10 mg/kg. A clear dose-dependent effect of MTD was observed in each of four separate measures: two locomotor measures (distance traveled in the arena and total time freezing) and two measures of vocal behavior (number of calls in 10 min and signal-to-noise ratio (SNR: defined as the amplitude of the vocalization signal above the background noise; or in other words, how “loud” the guinea pigs are, controlling for background noise) (Figure 2). To take better advantage of all data we also devised a composite outcome score for each animal based on z-transformation of all four measures (see Methods). This approach was modeled after clinical studies using scores constructed from multiple measurements or endpoints *(e*.*g*., *see [28])*. As shown in Figure 3, this approach not only *improved the continuity* of the dose-response (p<0.0001 for the linear fit), but also it *improved detection* of a significant effect at all three of the highest doses (note asterisks in Fig.3, and compare to Fig.2).

**Figure 2.**
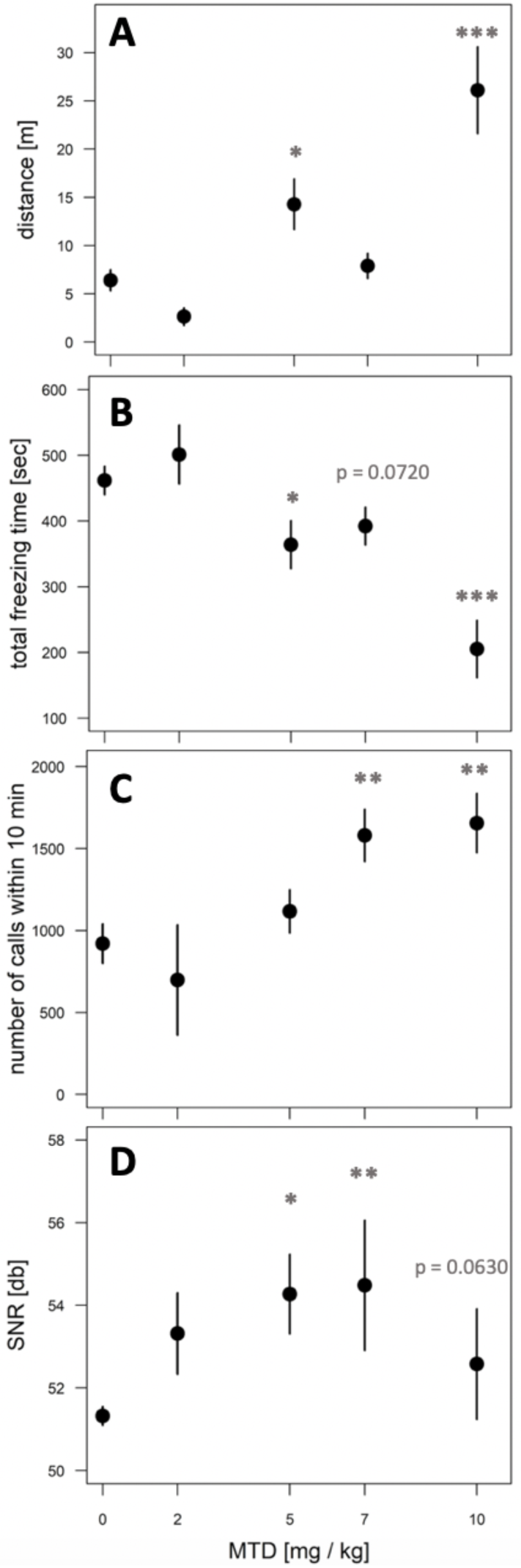
Dose-dependent effect of MTD on spontaneous withdrawal of newborns in the open-field. Pups at 48 hr after birth were placed in the open-field arena for 10 min and video recorded. A) Distance traveled in meters; B) total freezing time in seconds; C) total number of calls in 10 min; D) signal-to-noise ratio (SNR, or “loudness”) of calls. All behaviors show a significant overall effect of MTD by linear trend test: p < 0.001 for A, B; p < 0.01 for C; p < 0.05 for D. For individual comparisons to “0 MTD”, by Dunnett post-hoc test: *p < 0.05; **p < 0.01; ***p < 0.001. An average of 9 pups from 4 litters was used in each dosage group.

In contrast, after naloxone, many pups in all groups remained nearly completely immobile during the entire period of recording (Figure 4). The posture of immobile animals is reminiscent of the piloerection behavior described for the late or despair phase of maternal separation stress in many studies in guinea pigs and monkeys *(27)*. By contrast, other pups in some of the same groups showed classic naloxone-induced hyperlocomotion, and on rare occasions even jumping (not shown). Together the two contrasting behaviors of locomotion and immobility create a highly variable behavioral phenotype, and there is no overall significant effect of MTD in any measure after naloxone treatment (although there is a trend towards increasing locomotion distance with increasing MTD; F(4,42) = 2.38, p = 0.0674). In addition, if all naloxone data from the entire study are pooled, locomotion distance was significantly decreased in the 30 min test after naloxone compared to the 10 min test before naloxone (mean±SEM = 10.7±1.0 m before naloxone vs 6.5±1.2 m after naloxone, and F(1,202) = 7.06, p = 0.0085), which is not the expected result for a classic naloxone induced withdrawal response.

**Figure 3.**
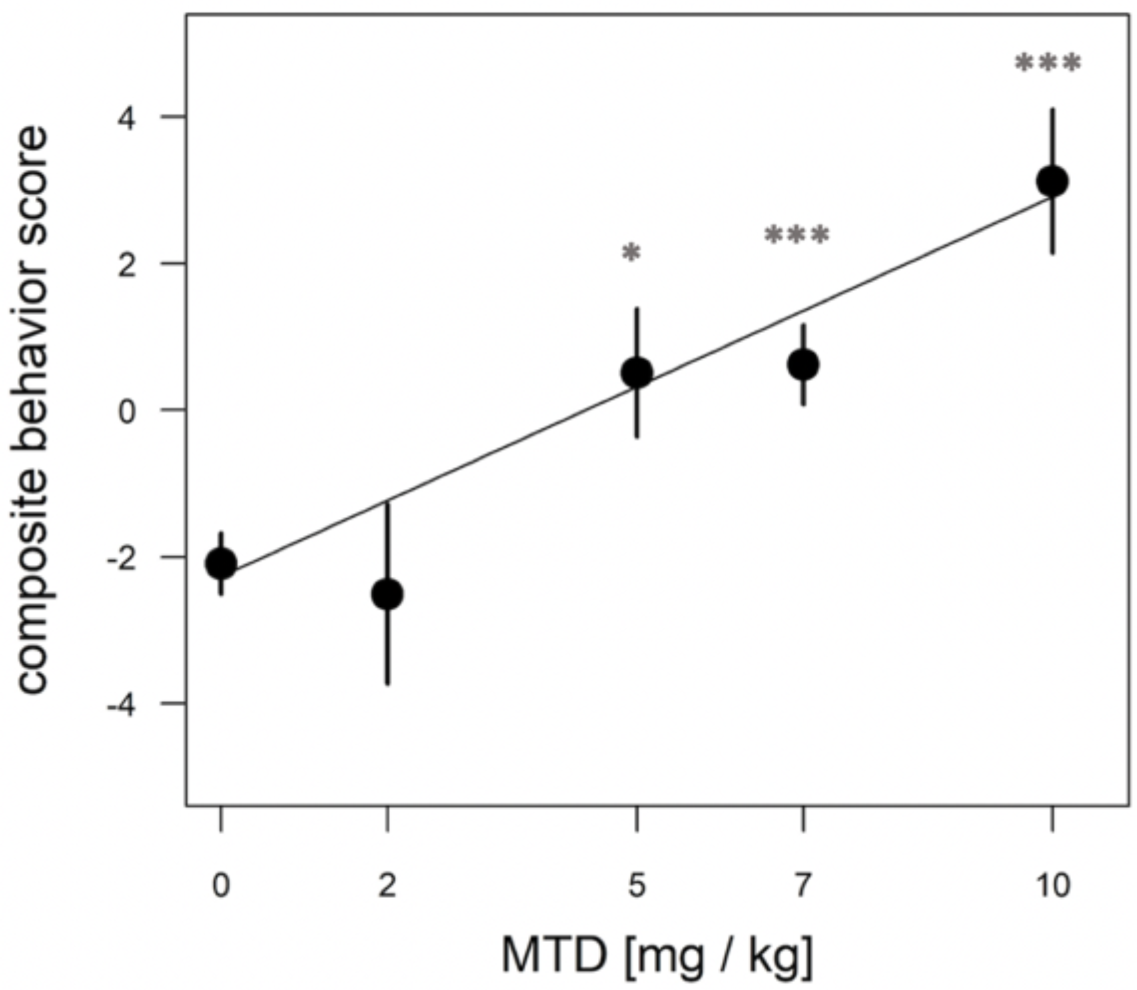
Dose-dependent effect of MTD using normalized composite outcome scores. Pups at 48 hr after birth were placed in the open-field arena for 10 min and video recorded. Data for distance traveled, total freezing time, number of calls in 10 min, and call SNR (“loudness”) were combined to generate composite outcome scores (see text for details). For overall effect of MTD p < 0.001 by linear trend test; * p < 0.05; *** p < 0.001, by Dunnett post-hoc test. An average of 9 pups from 4 litters was used in each dosage group.

**Figure 4.**
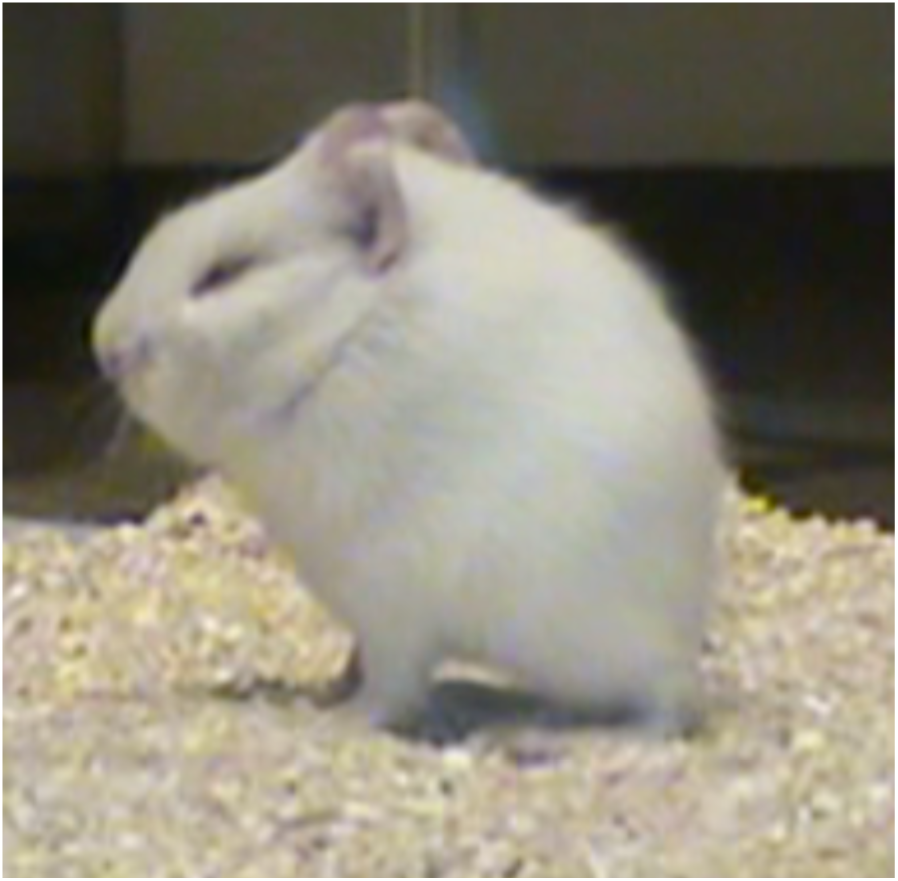
An example of immobility, piloerect posture and eye closure after naloxone induction of withdrawal in newborn guinea pigs exposed to prenatal MTD.

The above naloxone effect in newborns differs from adult and juvenile mice, which show robust naloxone-induced locomotion and jumping after several days of opioid exposure *(6,8,9)*. Here, we tested adult guinea pigs after 3 days of exposure to methadone (10 mg/kg/day). On day 4 they showed no significant increase in spontaneous locomotion relative to controls in a 10 min test in the open-field, but similar to mice, they show a robust increase in locomotion after naloxone injection; the converse of what was observed in newborns (Figure 5). Thus, newborn guinea pigs are unusual in not showing a significant naloxone-induced locomotor effect. This is in contrast to newborn rats exposed to a prenatal opioid, which display significant naloxone-induced increases in counts of limb and head movements *(15)*. In addition, the effect of prenatal MTD on spontaneous locomotion in newborn guinea pigs is an enhancement of a unique natural behavior for them, since newborn saline controls have a significant 6-fold higher level of locomotion than adult saline controls (Fig. 5A).

**Figure 5.**
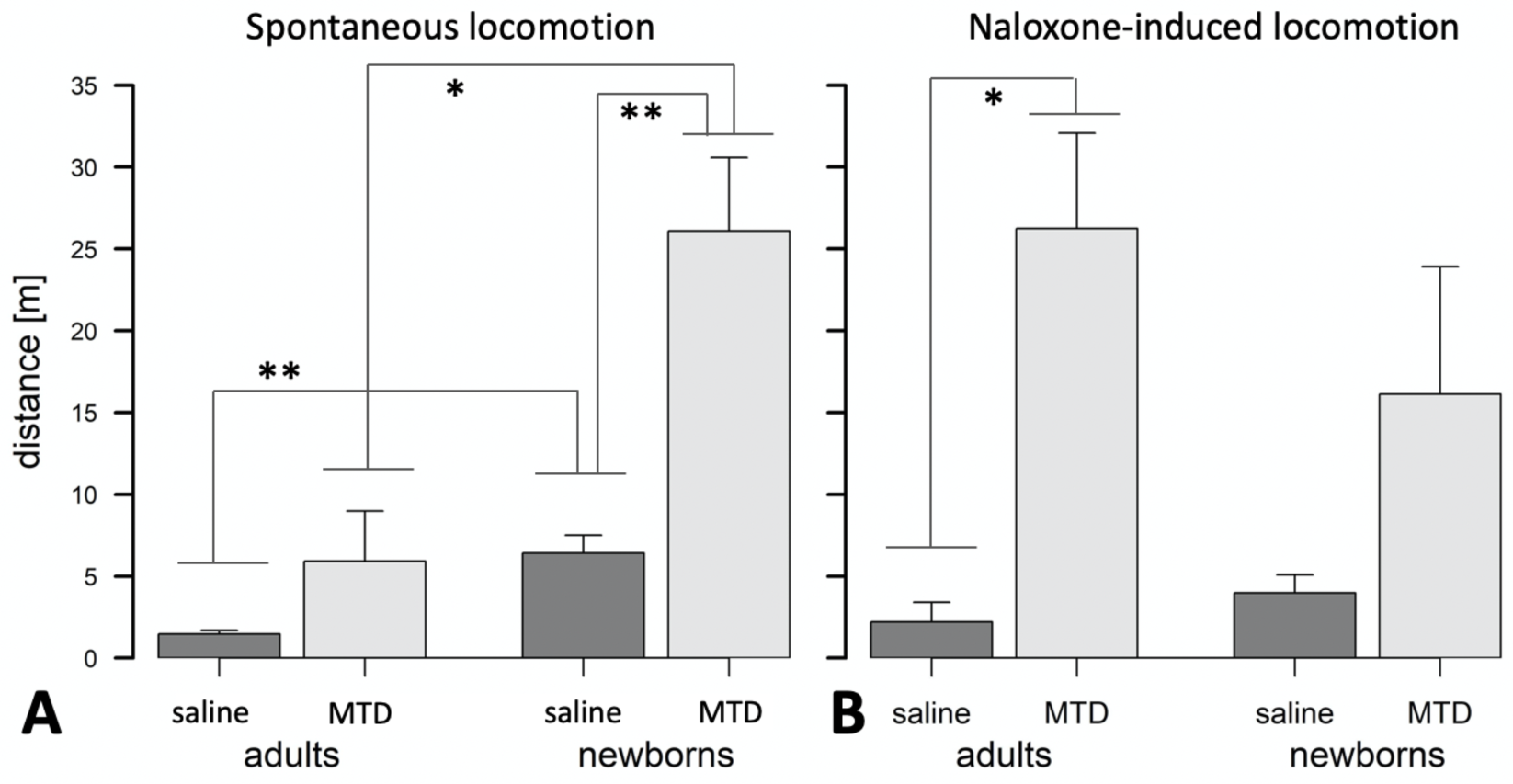
Spontaneous vs naloxone-induced withdrawal in adult and newborn guinea pigs. A) Adults treated with MTD show a trend, but no significant spontaneous withdrawal. Newborns with prenatal MTD show significant spontaneous withdrawal, and newborn controls are significantly more active than adult controls. B) Adults treated with MTD show significant naloxone-induced withdrawal, but newborns do not. A,B) MTD dose is 10 mg/kg; *p<0.05; **p<0.01, by t-test (with Bonferroni correction in A for multiple testing).

In sum, these observations support that the spontaneous increase in locomotion and calling observed in neonates is due to prenatal MTD’s effect on neonate-specific behaviors related to maternal separation, and not a general effect on locomotion and vocalization.

#### Naloxone-induced hypotonia

In association with the immobility behavior often observed in newborns after naloxone dosing there is an apparent sleep-related hypotonia (Supplemental Movie S2). The behavior always initiates in animals that are immobilized, and in an upright and slightly hunched posture. The behavior comes in two forms: either a brief wobble and the animal rapidly rights itself (an event), or the animal collapses on its side (a “gran mal” event). After collapse the animal may stay that way for a few seconds or even a few minutes, or right itself immediately. The behavior may be associated with yawning. Efforts to quantify this behavior by counting events plus gran mals, or based on the summed duration of all events and gran mals, showed no significant effect of MTD, either when the behavior was assessed on its own or when coupled to other behaviors such as locomotion using a composite score approach. Indeed, such events are even detectable in saline control animals receiving naloxone (and in all groups receiving MTD and 6BN together; see below). They were never observed in the 10 min videos before naloxone. In total, 74 out of 91 newborns (81%) in this study treated with either prenatal saline, prenatal MTD alone or with MTD plus 6BN (see below) showed at least one event, with an average of 3.8 events per animal, after treatment with naloxone. This naloxone-induced behavior is also observed in adults exposed to MTD, but it is much less prominent than for newborns (see below). In addition, the adult data support this behavior as a contributing factor in increased variability in locomotion and other measures after naloxone.

### 6BN prevents methadone’s effect on newborn withdrawal behaviors

We had reported that the partial exclusion of 6BN from the adult mouse brain was developmentally regulated and remained incomplete until 15 - 20 d after birth *(6)*. Using PD12-17 d juvenile mice as a model for fetal drug exposure in humans, we observed that 6BN had extreme potency to prevent morphine-induced dependence and withdrawal *(6)*. Also, in the same study we showed that brain AUC’s of 6BN were 6-fold higher in the fetal than the maternal brain at comparable blood levels owing to the immature BBB; whether this would result, as for juveniles, in high potency of 6BN to prevent dependence could not be tested, however, since mice at birth do not show evident withdrawal after prenatal opioid exposures *(14)*. Our goal is to test this in guinea pigs. However, because of slow placental transfer in pregnant guinea pigs, 6BN reaches only low absolute levels in fetal brain compared to mice, and roughly equal AUC’s in fetal versus maternal brain after a single 6BN dose (see Figure 1, this study). Therefore, based on a preferential delivery model whereby 6BN rapidly enters fetal brain but is relatively excluded from maternal brain, we would predict decreased 6BN potency for preventing neonatal withdrawal in pregnant guinea pigs due to slow placental transfer, as compared to juvenile mice, which have essentially no CNS barrier. Having established solid evidence of MTD effects on neonatal behaviors we set out to test this.

#### Effect of 6BN on maternal weight gain during pregnancy, and pup birthweight

For these studies most of the animals were tested with 5 mg/kg MTD and varying doses of 6BN. 5 mg/kg is at the midpoint of the dose-response curve (Figure 3). However, we also tested one cohort of animals at 7 mg/kg MTD and pooled the data with those at 5 mg/kg MTD (see Methods). First, we examined effects on pregnancy and birthweight. As shown in Table 6, MTD alone (“0 6BN”) yielded a trend for decreased maternal weight gain, consistent with the MTD effect described in Table 4, but 6BN did not significantly reverse this effect. However, 6BN *alone* at 0.3 mg/kg (no MTD; n = 2) significantly increased maternal weight gain even above the saline controls. There were no effects on litter size or pup survival. As shown in Table 7, MTD had no effect on pup birthweight consistent with data in Table 5. However, 6BN in conjunction with MTD (even at the lowest 6BN dose of 0.025 mg/kg), or 6BN alone (at 0.3 mg/kg), caused a significant ∼15% increase in pup birthweight above saline controls (F(4,149) = 5.48, p < 0.001, for overall effect of 6BN compared to saline controls, by ANOVA). In summary, this indicates an effect of 6BN to increase pup birthweight, and possibly maternal weight gain, independent of MTD treatment.

**Table 5.** Effect of methadone on pup birthweight

| MTD dose (mg/kg) | number of pups (n) | number of litters | birthweight (g) |
| --- | --- | --- | --- |
| 2 | 6 | 2 | 95.7±7.1* |
| 5 | 41 | 7 | 75.4±1.8 |
| 7 | 8 | 2 | 72.4±4.3 |
| 10 | 11 | 3 | 78.4±4.0 |
| 0 (saline ctl) | 28 | 5 | 75.5±3.1 |

all values ±SEM
no significant overall effect of MTD by ANOVA: F(4,89)=1.8671, p=0.1232
\*p < 0.05, for comparison to saline control, by Dunnetts posthoc test

**Table 6.**
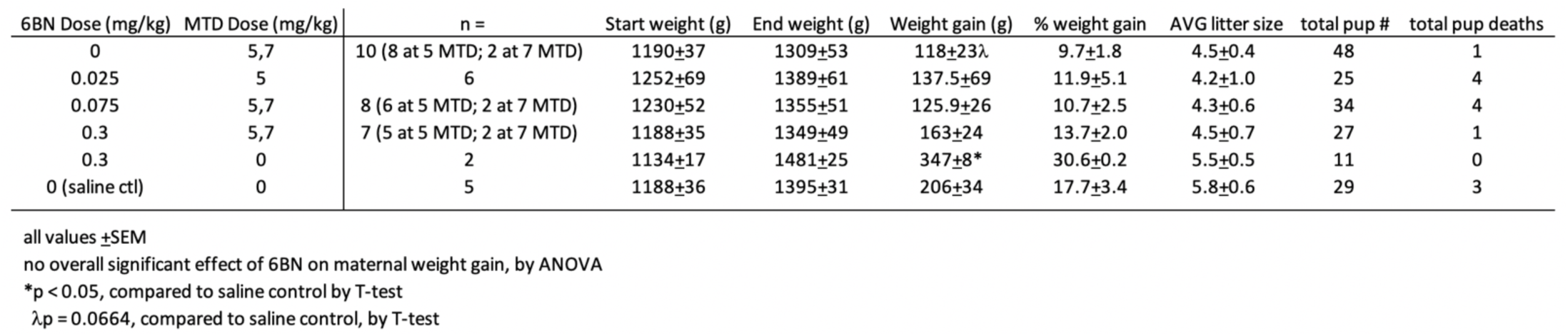
The effect of 6BN on maternal weight gain and litter size

**Table 7.**
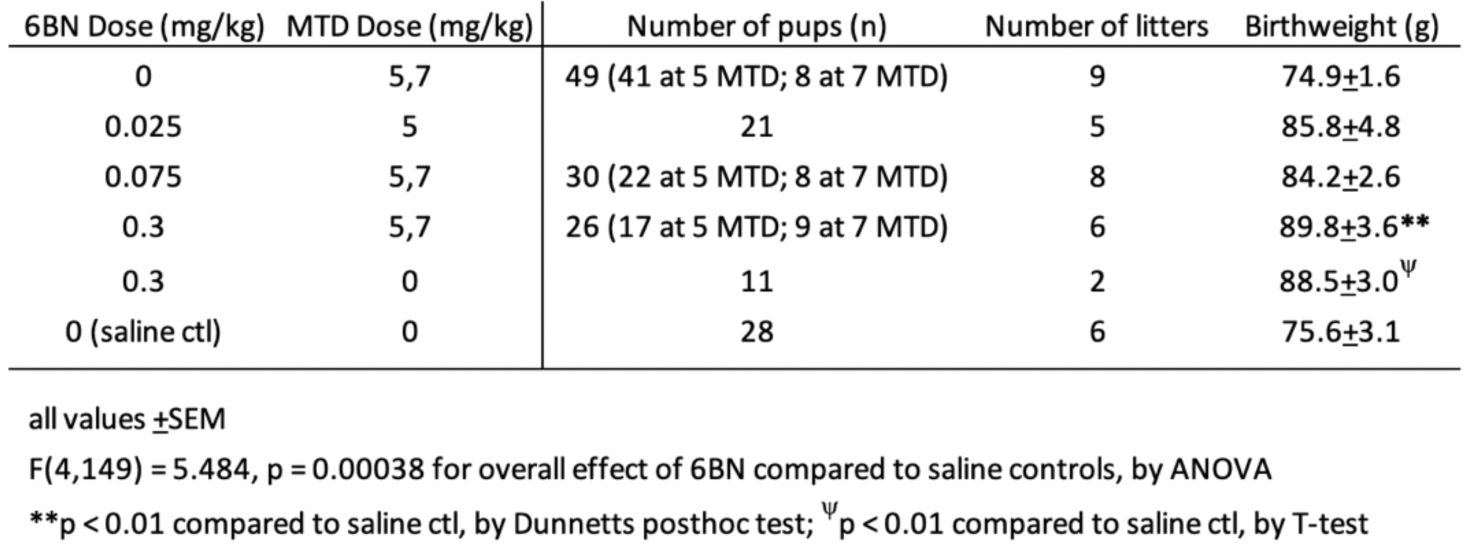
The effect of 6BN on pup birthweight

| 6BN Dose (mg/kg) | MTD Dose (mg/kg) | Number of pups (n) | Number of litters | Birthweight (g) |
| --- | --- | --- | --- | --- |
| 0 | 5,7 | 49 (41 at 5 MTD; 8 at 7 MTD) | 9 | 74.9±1.6 |
| 0.025 | 5 | 21 | 5 | 85.8±4.8 |
| 0.075 | 5,7 | 30 (22 at 5 MTD; 8 at 7 MTD) | 8 | 84.2±2.6 |
| 0.3 | 5,7 | 26 (17 at 5 MTD; 9 at 7 MTD) | 6 | 89.8±3.6** |
| 0.3 | 0 | 11 | 2 | 88.5±3.0<sup>ψ</sup> |
| 0 (saline ctl) | 0 | 28 | 6 | 75.6±3.1 |

all values ±SEM
F(4,149) = 5.484, p = 0.00038 for overall effect of 6BN compared to saline controls, by ANOVA
\*\*p < 0.01 compared to saline ctl, by Dunnetts posthoc test; <sup>ψ</sup>p < 0.01 compared to saline ctl, by T-test

#### Effect of 6BN on neonatal withdrawal behaviors

Next we tested whether 6BN could prevent neonatal withdrawal, at the doses of 5 & 7 mg/kg MTD as described above, applying the same four behavioral measures as used for the MTD dose-response. As shown in Figure 6, prenatal 6BN dosing combined with MTD had a significant overall effect on three of the four measures, reducing the effect of MTD to baseline at the highest 6BN dose used (0.3 mg/kg). One measure, total number of calls, did not show any significant effect. Using the composite outcome method described above, we observed a significant overall effect of 6BN on reducing MTD-induced spontaneous withdrawal (Figure 7A).

**Figure 6.**
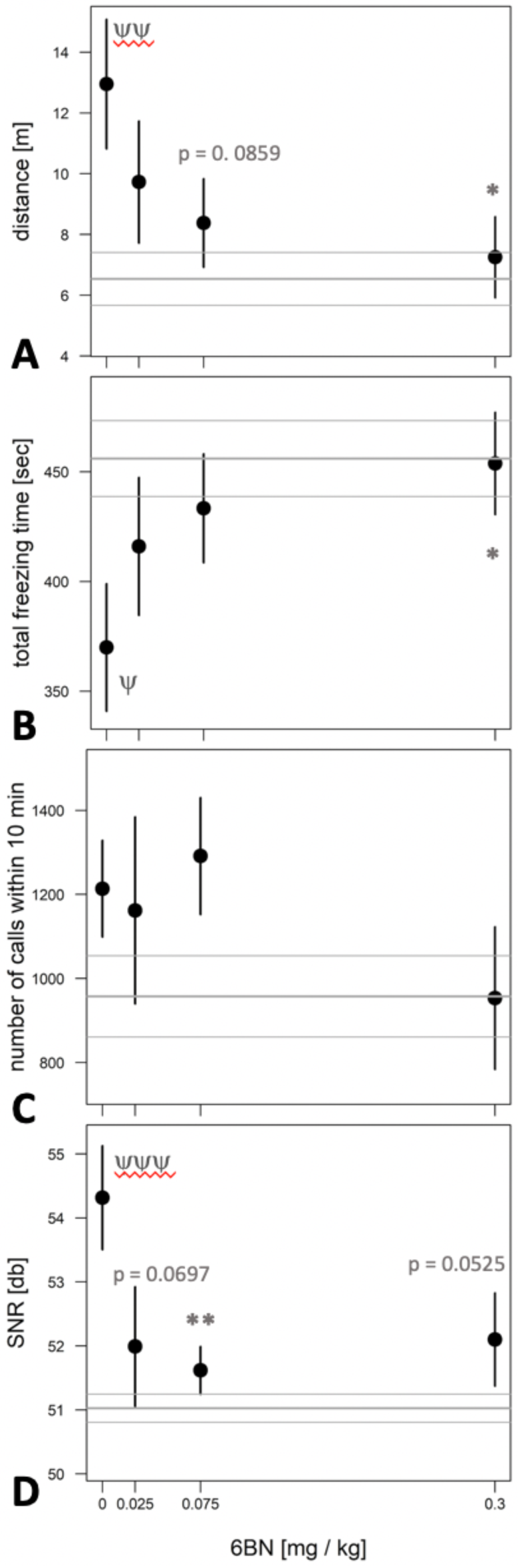
6BN prevents the effects of MTD on newborn withdrawal. Pups at 48 hr after birth were placed in the open-field arena for 10 min and video recorded. A) Distance traveled in meters; B) total freezing time in seconds; C) total number of calls in 10 min; D) signal-to-noise ratio (SNR, or “loudness”) of calls. Three behaviors show a significant overall effect of 6BN by linear trend test: p < 0.05 for A, B and D. For individual comparisons to “0 6BN”, by Dunnett post-hoc test: *p < 0.05; **p < 0.01. For comparison of “0 6BN” to saline control (grey horizontal lines, mean±SEM) by t-test: ^ψ^p < 0.05; ^ψψ^p < 0.01; ^ψψψ^p < 0.001. An average of 16 pups from 8 litters was used in each dosage group.

**Figure 7.**
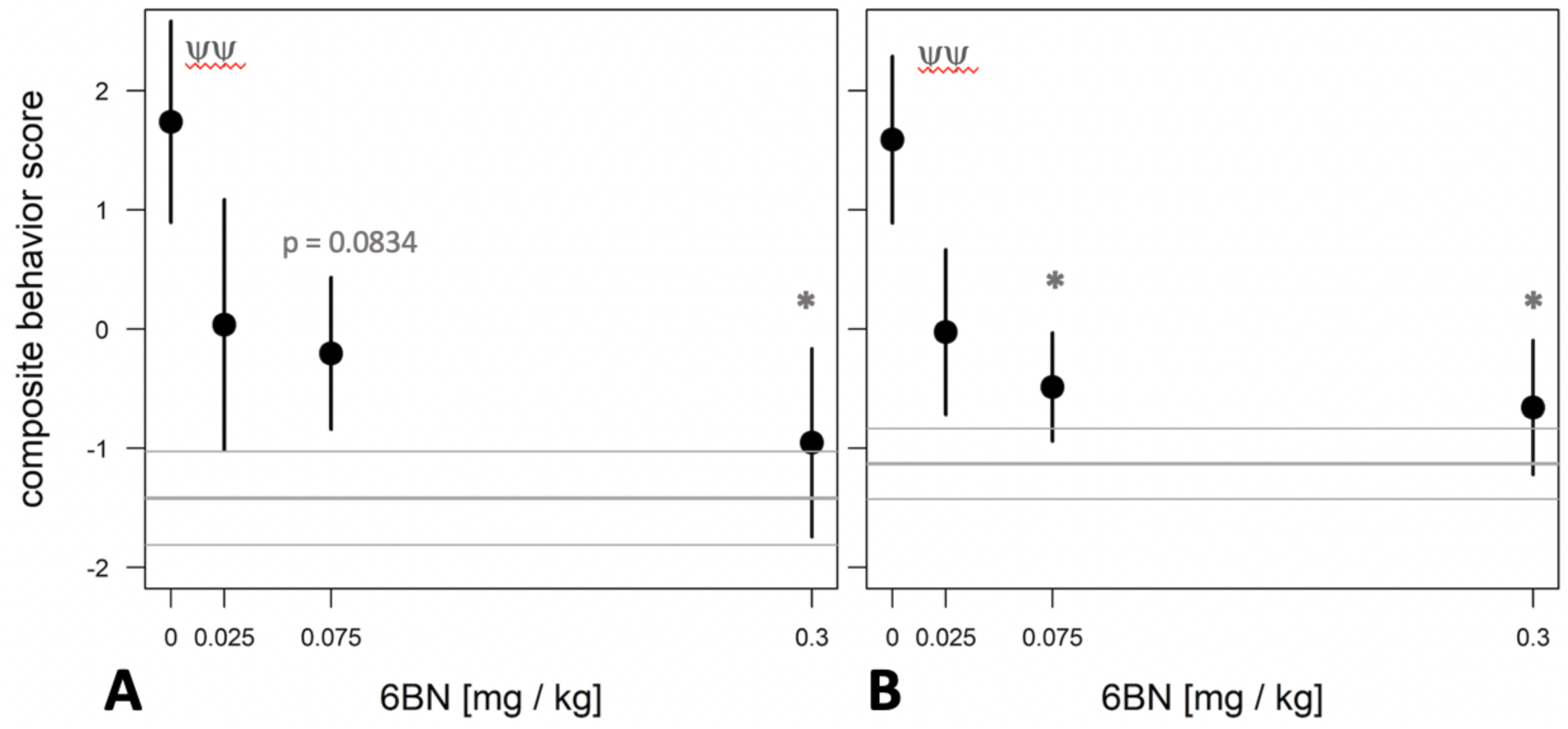
Dose-dependent effect of 6BN using normalized composite outcome scores. Pups at 48 hr after birth were placed in the open-field arena for 10 min and video recorded. A) Data for distance traveled, total freezing time, number of calls in 10 min, and call SNR (“loudness”) were merged to generate composite outcome scores (see text for details). B) Same as A, but without “number of calls” as a factor in composite score. In A,B: p < 0.05 for overall effect of 6BN by linear trend test; *p < 0.05 compared to “0 6BN”, by Dunnett post-hoc test; ^ψψ^p < 0.01 for “0 6BN” compared to saline control (grey lines, mean+SEM). An average of 16 pups from 8 litters was used in each dosage group.

However, the composite score did not increase statistical sensitivity as observed in the MTD dose-response study (Figure 3). This is likely due to the number of calls. To test this, we removed that measure, reanalyzed the composite score data, and observed an increased statistical sensitivity based on detection of a significant effect at a lower dose of 6BN (by individual comparisons) than that observed for any single measure alone (compare asterisks in Figs. 6 and 7B). These observations suggest that the two measures, number of calls and SNR, while increased by prenatal MTD exposure (Fig. 2), may be differentially affected by 6BN with number of calls requiring higher doses for its suppression.

When composite outcome data shown in Figure 7A were fitted with an emax model *(63)*, the following parameters were obtained: e0 (i.e., the behavioral score at 6BN dose = 0) = 1.73±0.70; emax (i.e, behavioral score at a maximal (infinite) 6BN dose) = - 2.724±1.29; and ID50 = 0.020±0.041. For the data shown in Figure 7B (i.e., a composite behavioral score without numcalls10), these parameters were estimated as follows: e0 = 1.59±0.54; emax = -2.35±0.91; and ID50 = 0.011±0.023. The range in ID50 values for 6BN indicated by this analysis is similar to what we reported in morphine-dependent juvenile mice (0.02 – 0.04 mg/kg; *(6)*). However, more dosing data at the lower and higher ends will be needed to calculate a more accurate ID50 by fitting a 4-parametric log curve, which awaits further studies. Nevertheless, it is quite clear that 6BN is as effective in pregnant guinea pigs to prevent neonatal opioid dependence as it is in juvenile mice, against two different agonists with distinctly different PK properties. This is in spite of slow placental transit of 6BN and its consequent low fetal brain levels in guinea pigs.

### MTD and 6BN effects on plasma cortisol levels

Previous studies have shown involvement of the hypothalamic-pituitary-adrenal (HPA) axis and neural-immune effects of maternal separation stress (MSS) *(25,29,30)*. Therefore, we tested the effect of MTD and 6BN on stress levels in newborns by measuring plasma cortisol. Plasma samples were collected from newborns 48 hr after birth that had been exposed to prenatal MTD with and without 6BN, as well as 1 hour of MSS just prior to sample collection. As shown in Table 8 animals exposed to prenatal MTD had 46% greater cortisol levels compared to saline injected controls (p < 0.05 by t-test). In addition, there was an overall significant effect of 6BN to prevent the MTD-induced increase in plasma cortisol (F(2,12) = 9.34; p = 0.0036 by ANOVA).

**Table 8.** 6BN prevents the increase in cortisol by prenatal MTD

| 6BN Treatment | Cortisol (ng/ml) |
| --- | --- |
| 0 | 1399 |
| 0.075 | 1094 |
| 0.3 | 859** |
| saline ctl | 958* |

-All pups treated with prenatal MTD (5 or 7 mg/kg) for 15 d with 6BN at the indicated dose, or with saline (no MTD, no 6BN); n = 5 pups from 3 litters per group = 3 pups at 5 mg/kg MTD and 2 pups at 7 mg/kg MTD in each group
-F(2,12) = 9.34, p = 0.0036 for overall effect of 6BN, by ANOVA
-\*p < 0.05, by T-test; \*\*p < 0.01 by Dunnett's posthoc analysis, all comparisons to 0 6BN condition

### MTD and 6BN effects in adult guinea pigs

Previous studies have reported low potency of 6BN in adult rodents and monkeys for blocking opioid antinociception and for inducing withdrawal in opioid-dependent animals, with an ID50 ranging from 1 to 10 mg/kg *(8-13)*. Therefore, the finding that 6BN *prevents* withdrawal at much lower doses in juvenile mice *(6)* or in newborn guinea pigs exposed to a prenatal opioid (this study) was unexpected. Here we tested whether 6BN could prevent withdrawal in adult guinea pigs, which, as described above in PK studies, show relatively poor brain entry of 6BN (Figure 1). As adult non-pregnant guinea pigs tolerate MTD better than the pregnant animals, we tested the ability of 6BN to prevent naloxone-induced withdrawal after 3 days of exposure to MTD at 10 mg/kg, twice the dose of MTD we used for the 6BN study in pregnant animals. In fact, when co-administered with MTD, 6BN reduced withdrawal by 71% at the dose of 0.03 mg/kg, the lowest dose showing a significant effect (Figure 8A). Using a dose-response emax model, the parameters for the exponential fitted curve are e0 = 19.6±3.62; emax = 4.5±5.3; and ED50 = 0.010±0.015. The mobility fitted for animals treated with the maximal dose (emax) was compared to the saline control (2.2±1.2) and cannot be distinguished from it (p = 0.86 by t-test), and thus prenatal 6BN effectively reduces the effect of MTD to levels of saline controls. The AIC (Akaike’s ‘An Information Criterion’, *(67)*) for this model is 166.77.

**Figure 8.**
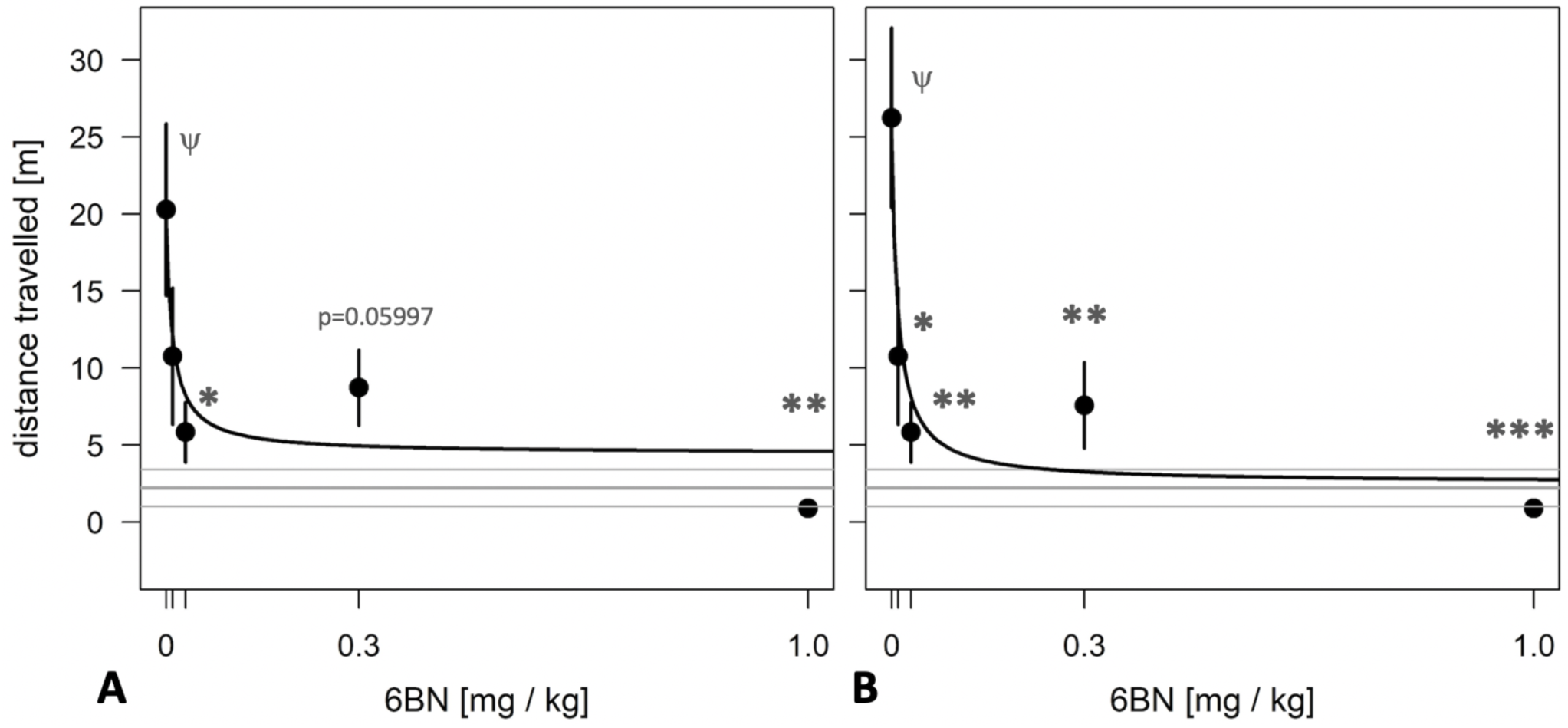
6BN has high potency to prevent opioid withdrawal in adult guinea pigs. Animals were treated for 3 d with one injection per day of 10 mg/kg MTD with varying doses of 6BN (0, 0.01, 0.03, 0.3, and 1 mg/kg). On day 4 animals were injected with naloxone and tested for 30 min in an open-field arena. A) 6BN delivered at low doses with MTD prevents naloxone-induced withdrawal in adult animals. p < 0.05 for overall effect of 6BN, by linear trend test. B) Same as A, but 3 animals with >15% of total arena-time in hypotonic posture were removed from dataset. p < 0.01 for overall effect of 6BN, by linear trend test. A,B) *p<0.05, **p<0.01, ***p<0.001, compared to “0 6BN” control, by Dunnett post-hoc test; ^ψ^p<0.05 compared to saline control (grey lines, mean±SEM), by t-test.

It should be noted that 8 of the 26 animals (31%) used in the adult study showed at least 1 bout of sleep-related hypotonia after naloxone treatment, with an overall average of 1 bout per animal for the adult dataset (see Supplemental Movie S3). Thus, this behavior is much less prominent than for newborns (see above). However, while 5 of these animals had only cursory bouts, spending less than 5% of their time in the arena in the prone hypotonic position, 3 of the animals spent greater than 15% of their time in this position. In addition, these 3 animals were either the highest or lowest in their respective groups for the measure of locomotion distance, suggesting an outsize influence on the data. To determine their effect on the overall curve shape and statistics we removed them from the dataset. As shown in Figure 8B, this allowed detection of a significant effect at all doses of 6BN, even at the lowest dose, 0.01 mg/kg. For this subset, the parameters for the exponential fit are e0 = 24.9±3.7; emax = 2.54±5.10; and ED50 = 0.010±0.009 (with upper limit of 0.036 mg/kg with 99% confidence), and again, the mobility fitted for the maximal dose (emax) cannot be distinguished from saline controls (p = 0.97 by t-test). The AIC for this model is 139.23. The reduced AIC (better exponential fit) in this subset compared to the complete dataset, and the increased sensitivity of detecting a significant effect at a lower dose in this subset, suggests that hypotonia is linked to increased data variability in the naloxone dataset. We did not observe a similar improvement in detection of 6BN effects in the newborn naloxone dataset using this approach (not shown), which we attribute to the much greater pervasiveness of this behavior in newborns.

## DISCUSSION

In this study we can draw two major conclusions. First, we have shown that prenatal MTD exposure aggravates classic maternal separation behaviors in newborn guinea pigs, including locomotor and vocal behaviors. It also significantly increases plasma cortisol in newborns, an indicator of an activated brain HPA axis. These results suggest a likely conduit for prenatal opioids and opioid cessation at birth to affect later-life brain development and behavior, through HPA activation, as shown in many studies of early life stress in diverse species from rodents to non-human primates *(31-34,42)*. It is also consistent with the recent suggestion that salivary cortisol in NOWS babies may be a suitable indicator of withdrawal severity *(35)*. In addition, animal studies, primarily in guinea pigs and monkeys, but also in rats, have suggested a role for endogenous opioids in the natural process of maternal attachment *(36,37,43)*. This suggests that cessation of an opioid at birth after chronic exposure in utero may result in an increased drive or craving for the opioid receptor stimulation provided by maternal contact; hence, increased calling and searching. This may explain the effectiveness of parent or surrogate rooming-in, breast feeding, and kangaroo care therapies for reducing ICU stay times and reducing the need for postnatal opioids in the clinic *(38-40)*.

The second major finding is that 6BN, when delivered together with the agonist, can prevent dependence-related behaviors in newborns exposed to extended periods of prenatal MTD - at 6BN doses unlikely to induce maternal or fetal withdrawal, or to interfere with opioid analgesia or use management. For example, the ID50 of 6BN for inducing withdrawal in opioid dependent rodents is in the range 1 – 10 mg/kg *(8,9)*, or 1, 1.3 or 2.4 mg/kg *(10-12)* for inhibition of opioid antinoception in rodents, or in the range of 0.3 – 1 mg/kg for interference of antinociception *and* induction of withdrawal in rhesus monkeys *(13)*. In addition, the potency of prenatally delivered 6BN to prevent withdrawal in newborn guinea pigs, with an estimated ID50 in the range of 0.01 – 0.02 mg/kg, is in a similar range as we observed in juvenile mice treated with morphine *(6)*. This high potency is in spite of the fact that MTD has a much longer half-life than 6BN, 12 hr for MTD in guinea pigs *(19)* versus ∼2 hr for 6BN (this study), and that 6BN has slow placental transit in guinea pigs (this study). Consistent with this result, we find that 6BN also has high potency to prevent withdrawal in adult guinea pigs, in which 6BN is relatively excluded from the CNS (also this study). A very similar observation has been made in adult mice prior to our study (*Z. Jim Wang, U. Illinois – Chicago; personal communication*).

From this evidence we conclude that barrier mechanisms at the placenta or BBB do not impede the potency of 6BN in preventing neonatal or adult dependence, and other mechanisms may be in play to explain its relatively low potency to induce withdrawal in dependent animals or to interfere in opioid analgesia. One possibility is a greater role of the peripheral nervous system in driving withdrawal behaviors than is currently appreciated. For example, recent studies have shown that auricular stimulation of cranial nerves reduces withdrawal symptoms, and may act by reducing sympathetic activity (“fight or flight”) that is increased during withdrawal, favoring parasympathetic predominance and reducing physical withdrawal symptoms *(41)*. Preferential blockade of opioid actions on cranial nerves by 6BN could maintain normal function of these peripheral neurons so that when the opioid ceases there is reduced sympathetic imbalance. Alternatively, 6BN may interact in novel ways with the opioid receptor, possibly binding with high affinity to a distinct receptor conformation involved in the development of dependence. For example, Jeske *(53)* has proposed a receptor model with distinct peripheral and central mu opioid receptor forms that could in part account for the observed peripheral selectivity of 6BN, even in rhesus monkeys where it readily penetrates the adult’s brain (J. Oberdick, unpublished results; see also *[68]*). It appears that our current understanding of the molecular pharmacology of opioid receptors is incomplete – requiring a new approach to account for the high potency of 6BN in selectively preventing opioid withdrawal behavior in dependent animals, as observed here and in mice *(6,68)*.

Several novel features of the guinea pig model are worthy of discussion. First, the behavior of guinea pig newborns enables a focus on maternal separation stress immediately after birth. This two-stage process mimics what is observed in non-human primates *(25)*, and since it clearly involves the HPA axis, neuroinflammatory mechanisms are immediately suggested *(29,30)*. In contrast, rats at birth show only modest behaviors, which, while statistically significant, are not immediately relatable to specific CNS neuronal pathways *(15,16)*. Not until well after birth, in the period PD7 – PD10, do rat pups show evidence of a clear maternal separation stress response. As in guinea pig newborns this response in PD7-10 rats appears to be dependent on endogenous opioids *(43)*, and it is aggravated by daily opioid treatments for the first postnatal week with cessation on PD7 *(44)*. While these 7-day old rat pups may serve as a reasonable model for some features of NOWS, they lack the temporal feature of withdrawal that is initiated due to opioid cessation at birth.

The second novel feature in newborn guinea pigs is the close apposition of two behaviors, locomotion and vocalization, that are at the heart of the maternal separation stress response. These two behaviors are developmentally coordinated in a manner linked to arousal, and thus their coordination is highly influenced by environment *(45)*. Thus, multiple but distinct measures of each of these two behaviors may be useful tools for dissecting neuronal pathways. In particular, SNR (or “loudness”) may be a vocalization feature that most closely relates to animal stress, and the magnitude of increase in stress vocalization that we observed when comparing saline controls to prenatal opioid exposed newborns, ∼4 db, is equivalent to the magnitude of change observed during castration of unanesthetized newborn pigs in a farm setting *(46)*. This comparison may highlight that newborns exposed to prenatal opioids are under considerable stress. Also, while number of calls is a measure most often used to assess changes in vocalization in mouse reverse genetics studies (e.g., *(47)*), the fact that this measure is not significantly suppressed by 6BN in newborn guinea pigs exposed to prenatal opioids, while plasma cortisol levels are suppressed, may suggest that counts of number of calls are not as relevant to stress as is SNR.

A third novel feature of the guinea pig as observed here is the sleep-related hypotonia induced by naloxone. The behavior is often accompanied by yawning, which supports that it is sleep-related. The observation that it is much more prominent in newborns than adults may be due to differences in physiology, but the fact that it is observed even in pups that received prenatal saline instead of MTD may suggest a strong influence of endogenous opioids, which as described above likely drives natural infant-maternal attachment *(36,37,43)*. In adults it appears that the relatively less prevalent version of this naloxone-induced behavior is specific to animals dependent on exogenous opioids, but more studies are needed. Disrupted sleep patterns is a well-described feature of NOWS in the clinic *(64,65)*, and therefore circadian activity studies may be of value in the future to study the effects on sleep and wakefulness in guinea pig newborns exposed to prenatal opioids.

The final feature worthy of discussion is the ability to combine these behaviors into a single composite score, at the level of each individual animal. This feature may allow for both increased efficiency of animal usage in terms of statistical detection, but also, by providing a more robust read-out of withdrawal, it may allow better understanding of the relationship between neonatal withdrawal severity and developmental consequences in later-life.

In conclusion, here we report a robust behavioral model of NOWS in pregnant guinea pigs, and we take a first step towards testing 6BN as a preventive therapy for NOWS, with a focus on newborn outcomes. Further studies are needed to determine the limits of 6BN potency, such as at higher doses of MTD, or with other commonly used opioids in the clinic such as buprenorphine. In addition, further studies are needed to specifically examine, in adult and pregnant guinea pigs, the levels of 6BN needed to either induce withdrawal in opioid dependent animals or to interfere with opioid antinociception. 6BN has been universally shown to have low potency for these actions in both adult rodents and monkeys, and it will be important to contrast this with low levels needed to prevent neonatal withdrawal in a single animal model. In addition, it remains to be seen whether initiation of MTD and 6BN combined therapy in pregnant animals that are already MTD-dependent, a condition likely to apply in the clinic, causes any untoward maternal or fetal effects related to withdrawal. Thus, in future studies we will focus much more on effects of opioids and 6BN on maternal outcomes.

## Acknowledgements

This work was supported by NIH R21-HD092011 to J.O. and NIH R44-DA045414 (SBIR) to W.S. Support for analysis of locomotor behavior was also provided by a P30 Core grant (NINDS P30-NS045758). Opioid agonists and antagonists were provided by the NIDA Drug Supply Program.

## AUTHORSHIP CONTRIBUTIONS

Performing experiments: J.O., A.S., S.A., V.A.M. guinea pig husbandry, tissue collection, and behavior; K.H, K.D mass spectrometry

Experimental design: J.O., K.L.B., W.S., and M.A.P.

Scoring of behavior by analysis of video recordings: A.R.L. (vocalization) and J.F. (locomotion)

Statistical analysis: M.C. and M.A.P. (pharmacokinetics) and K.S. (behavior)

Manuscript writing and revision, and financial support: J.O., S.A., A.S., K.S., A.R.L., K.L.B., K.O., M.A.P., W.S.

